# Adipose depot-specific upregulation of Ucp1 or mitochondrial oxidative complex proteins are early consequences of genetic insulin reduction in mice

**DOI:** 10.1101/2020.03.31.018432

**Authors:** Jose Diego Botezelli, Peter Overby, Lorenzo Lindo, Su Wang, Obélia Haïda, Gareth E. Lim, Nicole M. Templeman, Jose Rodrigo Pauli, James D. Johnson

**Affiliations:** Department of Cellular Physiological Sciences, Diabetes Research group, Life Sciences Institute, University of British Columbia, Vancouver, BC, Canada; Laboratory of Molecular Biology of Exercise (LaBMEx), Faculty of Applied Sciences, University of Campinas, Limeira, São Paulo, Brazil; Cardiometabolic axis, Centre de recherche du Centre hospitalier de l’Université de Montréal, Université of Montréal, Montréal, QC, Canada; Princeton University, Princeton, New Jersey, USA

**Keywords:** hyperinsulinemia, adipose tissue, obesity, uncoupling protein 1, oxidative phosphorylation

## Abstract

Hyperinsulinemia plays a causal role in adipose tissue expansion. Mice with reduced insulin have increased energy expenditure, but the mechanisms remained unclear. Here we investigated the effects of genetically reducing insulin production on uncoupling and oxidative mitochondrial proteins in liver, skeletal muscle, white adipose tissue (WAT), and brown adipose tissue (BAT). Male *Ins1*^+/+^ or *Ins1*^+/-^ littermates were fed either a low-fat diet (LFD) or a high-fat diet (HFD) for 4 weeks, starting at 8 weeks of age. Replicating our previous observations, HFD increased fasting hyperinsulinemia, and *Ins1*^+/-^ mice had significantly lower circulating insulin compared with *Ins1*^+/+^ littermates. Fasting glucose and body weight were not different between genotypes. We did not observe significant differences in liver in skeletal muscle. In mesenteric WAT, *Ins1*^+/-^ mice had reduced Ndufb8 and Sdhb. Ucp1 was increased in the context of the HFD, and HFD alone had a dramatic inhibitory effect on Pparg abundance. In inguinal WAT, *Ins1*^+/-^ mice exhibited significant increases in oxidative complex proteins, independent of diet, without affecting Ucp1, Pparg, or Prdm16:Pparg association. In BAT, lowered insulin increased Sdhb protein levels that had been reduced by HFD. Ucp1 protein, Prdm16:Pparg association, and Sirt3 abundance were all increased in the absence of diet-induced hyperinsulinemia. Our data show that reducing insulin upregulates oxidative proteins in inguinal WAT without affecting Ucp1, while in mesenteric WAT and BAT, reducing insulin upregulates Ucp1 in the context of HFD. Preventing hyperinsulinemia has early depot-specific effects on adipose tissue metabolism and help explain the increased energy expenditure previously reported in *Ins1*^+/-^ mice.

## Introduction

Insulin is secreted from pancreatic β-cells in response to nutrient intake to regulate anabolism in multiple tissues. In addition to glucose homeostasis, insulin is a potent regulator of lipid homeostasis (42). Insulin signaling is required for glucose and lipid uptake into fat, as well as for the development and hypertrophy of adipocytes. Indeed, mice lacking insulin receptors in adipose tissue are protected against high fat diet and glucose intolerance (3, 34). Chronically elevated insulin levels are associated with obesity in human populations and pre-clinical models (1, 6, 27, 31, 39). Pharmacological inhibition of insulin secretion with diazoxide or octreotide can cause weight loss (2, 40). However, caveats related to the non-specificity of these drugs beyond insulin secretion meant that the causality of insulin in obesity had not been firmly established.

Using mice with specific genetic reductions in insulin secretion, we and others have shown that hyperinsulinemia contributes causally to diet-induced obesity, age-dependent insulin resistance, fatty liver, and accelerated aging, across both constitutive and inducible models and in both sexes (7, 22, 25, 37, 43). Mechanistically, suppression of hyperinsulinemia increased energy expenditure in young male mice prior to the divergence in body weight at ∼24 weeks of age (22). In tissues isolated from those mice at 1 year of age, we observed up-regulation of *Ucp1* and *Pparg* mRNA and protein in white adipose tissue (WAT)(22), but it is not known whether these molecular alterations were causes or consequences of changes in body weight. Collectively, our previous observations point to critical roles for insulin in mitochondria function in the context of obesity, however the molecular mechanisms underlying the increased energy expenditure in mice with reduced insulin production had not been fully elucidated.

In the present study, we investigated the effects of genetically reducing insulin production on the abundance of oxidative mitochondrial protein complexes and other proteins known to affect energy expenditure in different tissues prior to differences in body weight. We found that mice with reduced insulin gene dosage show differential modulation of mitochondrial oxidative complex protein abundance among mesenteric WAT, inguinal WAT, and brown adipose tissue (BAT) after just 4 weeks of diet intervention, and no consistent changes in these proteins were detected in liver or muscle. The increased *in vivo* energy expenditure we have previously measured in mice with reduced insulin production is therefore associated with upregulation of mitochondrial oxidative complex proteins in inguinal WAT, but an increase in Ucp1 protein in mesenteric WAT and BAT. These results shed light on the effects of hyperinsulinemia on different classes of adipocytes.

## Materials and Methods

### Animal Husbandry

Animal procedures were approved by the University of British Columbia Animal care Committee (A16-0022). All mice were maintained on an *Ins2* null background to prevent genetic compensation, as described previously (19) (21). Animals were housed at specific pathogen-free Centre for Disease Modeling (CDM) during the entire experiment. Eight week old control *Ins1*^+/+^;*Ins2*^-/-^ mice and experimental *Ins1*^+/-^;*Ins2*^-/-^ mice were randomly distributed into groups *ad libitum* fed either low fat diet (Research Diets D12450B, 20% protein 10% fat, 70% carbohydrate content, energy density 3.82Kcal/g, Brunswick, NJ, USA) or high fat diet (Research Diets D12492, 20% protein, 60% fat, 20% carbohydrate content, energy density 5.21Kcal/g, Brunswick, NJ, US) for 4 weeks. Mice were housed 12/12h light/dark cycle in the UBC Centre for Disease Modeling (CDM) under SPF conditions at the temperature of 21°C. Animals had body weight and glucose levels recorded every weekly.

**Table 1.** Diet nutrient description.

| **Class Description** | **Ingredient** | **Low Fat Diet (g)** | **High Fat Diet (g)** |
| --- | --- | --- | --- |
| **Protein** | Casein | 200.00 | 200.00 |
| **Amino Acid** | Cysteine L | 3.00 | 3.00 |
| **Carbohydrate** | Sucrose | 354.00 | 72.80 |
| **Carbohydrate** | Starch | 315.00 | - |
| **Carbohydrate** | Lodex 10 | 35.00 | 125.00 |
| **Fiber** | Solka Floc FCC200 | 50.00 | 50.00 |
| **Fat** | Soybean Oil, USP | 25.00 | 25.00 |
| **Fat** | Lard | 20.00 | 245.00 |
| **Mineral** | RD-96 Mineral Mix | 50.00 | 50.00 |
| **Vitamin** | Choline Bitartrate | 2.00 | 2.00 |
| **Vitamin** | AIN-76A Vitamin Mix | 1.00 | 1.00 |
| **Total:** |  | 1055.05g | 773.85g |

### Glucose and Insulin Measurements

Mice were fasted for 4h in clean cages during the light period (8:00-12:00 am) prior to assessing blood glucose using a OneTouch Ultra2 glucose meter (LifeScan Canada Ltd, Burnaby, BC, Canada). Plasma insulin was measured with a mouse insulin ELISA kit (Alpco Diagnostics, Salem, NH, USA) according to the manufacturer’s protocol.

### Tissue Extraction and Oxygen Consumption Measurements

Mice were fasted for 4h in clean cages (8:00-12:00 am) before the euthanasia. The liver central lobe, gastrocnemius muscle, mesenteric WAT, inguinal WAT, interscapular BAT, were dissected, isolated and sliced in parts weighing 5-10 mg using a scalpel and a petri dish on ice. Fat tissues were isolated and minced into ∼2-1 mg sections. All samples were washed in 1xPBS to remove hair and blood and transferred to fresh wash media (eagle’s minimal essential medium (DMEM, no glucose) (Gibco^®^) supplemented with 25 × 10-3 M glucose and 25 × 10-3M 4-(2-hydroxyethyl)-1-piperazinemethanesulfonic acid (HEPES buffer). A pair of sterile forceps were used to position 1 section of tissue into each well in the Seahorse XF96 spheroid plates containing seahorse assay media (DMEM, 10mM glucose). The Seahorse XF96 (Seahorse Bioscience, North Billerica, MA) was used to measure basal OCR. This system measures changes in oxygen concentration in a small amount of media that is isolated above the tissue. Basal OCR was normalized to dry tissue weight. Samples were stored at −80°C.

### Co-Immunoprecipitation and Western Blot Analysis

Fractions (5-10 mg) of liver, gastrocnemius muscle, mesenteric WAT, inguinal WAT, and interscapular BAT were placed in ice-cold RIPA buffer and lysated in TissueLyser (TissueLyser II, Qiagen, Vancouver, Canada). The samples were sonicated and centrifuged at 14,000 g for 20 minutes at 4°C. The supernatant protein concentration was determined using a BCA assay kit (Pierce BCA Protein Assay Kit, #23227, Thermo Fischer Scientific, Whalam, USA). Aliquots of 30 μg protein were mixed with Laemmli loading buffer, electrophoresed in 10% SDS-PAGE gels, and transferred in a wet tank to PVDF membranes (Immun-Blot PVDF Membrane, #1620177, Bio-Rad, CA, USA). Membranes were blocked in Tris-Buffered saline containing iBlock (iBlock™, #T2015, Applied Biosystems, California, USA). Blocked membranes were incubated at 4°C overnight in iBlock-Tris solution containing primary antibody. A second incubation containing a horseradish peroxidase-conjugated secondary antibody was performed at room temperature for two hours. Membranes were then washed using a commercial HRP Substrate solution (Immobilion Forte Western HRP, #WBLUF0100, Millipore Sigma, Oakville, Canada). After protein quantification, membranes were washed in restore/stripping buffer (Restore™ Western Blot Stripping Buffer, #21059, ThermoFisher Scientific, Waltham, USA) and incubated in another primary antibody solution, or Actinb as a loading control.

For co-immunoprecipitation experiments, 100 μg of protein was incubated overnight in the primary antibody Prmd16 (anti-Prdm16, #ab106410, Abcam, Whitby, Canada) followed by a 4-h incubation in Protein A magnetic beads (Pierce™ Protein A Magnetic Beads, #88845, ThermoFisher Scientific, Waltham, USA) and isolated using a magnetic rack. The isolated immunoprecipitated pellet was washed and further used for a western blotting assay. Protein expression density was accessed using the software Photoshop CC (Adobe^®^, San Jose, California, USA).

The immunoblotting was performed using antibodies for mitochondrial complexes (Total OXPHOS Rodent WB, #ab110413, Abcam, Whitby, Canada), Pparg (Anti-Pparg, #ABN1445, Millipore Sigma, Oakville, Canada), Ucp1 (Anti Ucp1, #PA5-29575, ThermoFisher Scientific, Waltham, USA), Prdm16 (anti-Prdm16, #ab106410, Whitby, Canada) and Actb (anti-beta-actin, #AC-15, Novus Biologicals, Oakville, Canada).

### Sirtuin Activity

For Sirt1 activity, a nuclear extraction kit (Nuclear Extraction Kit #ab113473, Abcam, Whitby, Ontario, Canada) was used to extract and isolate the nuclear protein fractions of a 5-10 mg sample of BAT. Later, we used the bicinchoninic acid method (Pierce™ BCA Protein Assay Kit #23225, ThermoScientific™, Burnaby, Canada) to measure the protein concentration of the homogenate. Samples were then, diluted to a final concentration of 2 µg/µL in nuclei extract buffer. Nuclear sirtuin 1 content and activity was assessed using 10 µg (5 μL) of isolated nuclear protein fractions through a fluorometric assay (Sirt1 Activity Assay Kit, #ab156065, Abcam, Whitby, Ontario, Canada). Another set of samples weighing 5-10 mg were homogenized in RIPA buffer (150 mM sodium chloride, 1.0% Triton x-100, 0.5% sodium deoxycholate, 0.1% SDS and 50 mM Tris) in a tissue lyser (TissueLyser II, Qiagen, Vancouver, CA) then sonicated for 10 minutes in a Q500 sonicator (QSonica, Newtown, USA) using 30s pulse intervals with 80% amplitude, before being centrifuged at 14,000 g for 20 minutes. The supernatant used for protein concentration determination via the BCA method. The samples were diluted in RIPA buffer to a final concentration of 2 μg/µL. 10 μg (5 μL) of homogenate was used to perform the Sirt3 concentration and activity using a fluorometric kit (SIRT3 Activity Assay Kit, #ab156067, Abcam, Whitby, Canada) according to the manufacturer’s instructions.

### 3T3L1 Adipocyte Cell Line Culture

3T3L1 pre-adipocytes (ZenBio; Research Triangle Park, NC) were maintained in 25 mM glucose DMEM, supplemented with 10% newborn calf serum and 1% penicillin/streptomycin. 3T3L1 cells were differentiated by allowing cells to reach confluence, followed by a cocktail (MDI) 500 μM IBMX, and 500 nM dexamethasone in differentiation media containing 25 mM glucose DMEM, 10% fetal bovine serum and insulin as indicated. Two days following induction, cells were maintained in differentiation media with insulin, and after 7 or 14 days, cells collected for RNA or protein or stained with Oil Red-O to assess adipogenesis (optical density).

### Statistical Analysis

Glucose concentration and body weight time course data were analysed using R Studio 3.4.1. Significance level of adjusted p < 0.05 was used. Both sets were analysed using linear regression modeling. Linear mixed effect models (R package – lme4) were fitted using restricted maximum likelihood. Predictor variables were included as fixed effects and sample IDs were included as random effects. Mixed effect modeling was used to account for repeated sample measurements and missing data. Heteroscedasticity and normality of residuals were analyzed used Levene’s test and the Shapiro–Wilk test, respectively. Predictor variables: genotype, diet, and time, were treated as categorical, categorical, and continuous factors, respectively. The outcome variables, glucose concentration or body weight, were treated as a continuous factor. Multiple comparison p-values were adjusted using the Tukey method. For the other parameters (Western blotting and activity assays), statistical data was performed using Statistica 12.0 software (Dell Software, Round Rock, USA). Two-way ANOVA was used to address the factors of diet, genotype and interaction between both factors. Results generated by the ANOVA were subjected to a Bonferroni post-hoc test. The significance level was set at *p* < 0.05. Graphpad Prism 7 software (Graphpad Software, San Diego, USA) was used to generate all graphs and images.

## Results

### Effects of diet and Ins1 gene dosage on circulating insulin, glucose, and body weight

The purpose of this study was to define the molecular mechanisms, at the protein level, of the increased energy expenditure observed in our previously published investigation on the role of hyperinsulinemia on weight gain (22). We have previously shown that male *Ins1*^+/*-*^ mice have lower circulating insulin when compared with *Ins1*^+/+^ littermate control mice at multiple time points (22). In the present study, conducted in a different animal facility, but under the same temperature housing conditions, we randomized male *Ins1*^+/+^ or *Ins1*^+/*-*^ mice into 2 diet groups and sought to replicate our previous observations (**Figure 1A**). Indeed, mice fed a HFD for 4 weeks had increased circulating insulin (LFD vs HFD; p=0.03). Deletion of one *Ins1* allele reduced this hyperinsulinemia (*Ins1*^+/+^ versus *Ins1*^+/*-*^; p=0.05) **(Figure 1B,C)**, confirming our previous findings (22). At the end of 4 weeks, we did not find significant differences in fasting glucose or body weight between any of the experimental groups (**Figure 1D,E**). These findings in 12 week-old mice with 4 weeks of diet intervention are consistent with our previous data illustrating that mice with reduced insulin gene dosage did not have robust differences in body weight prior to ∼24 weeks of age (22). Therefore, this relatively acute model provided an opportunity to examine, prior to differences in body weight or blood glucose, the protein abundances of potential upstream mediators of energy expenditure in key tissues that could affect the increased energy expenditure in this model (22).

**Figure 1.**
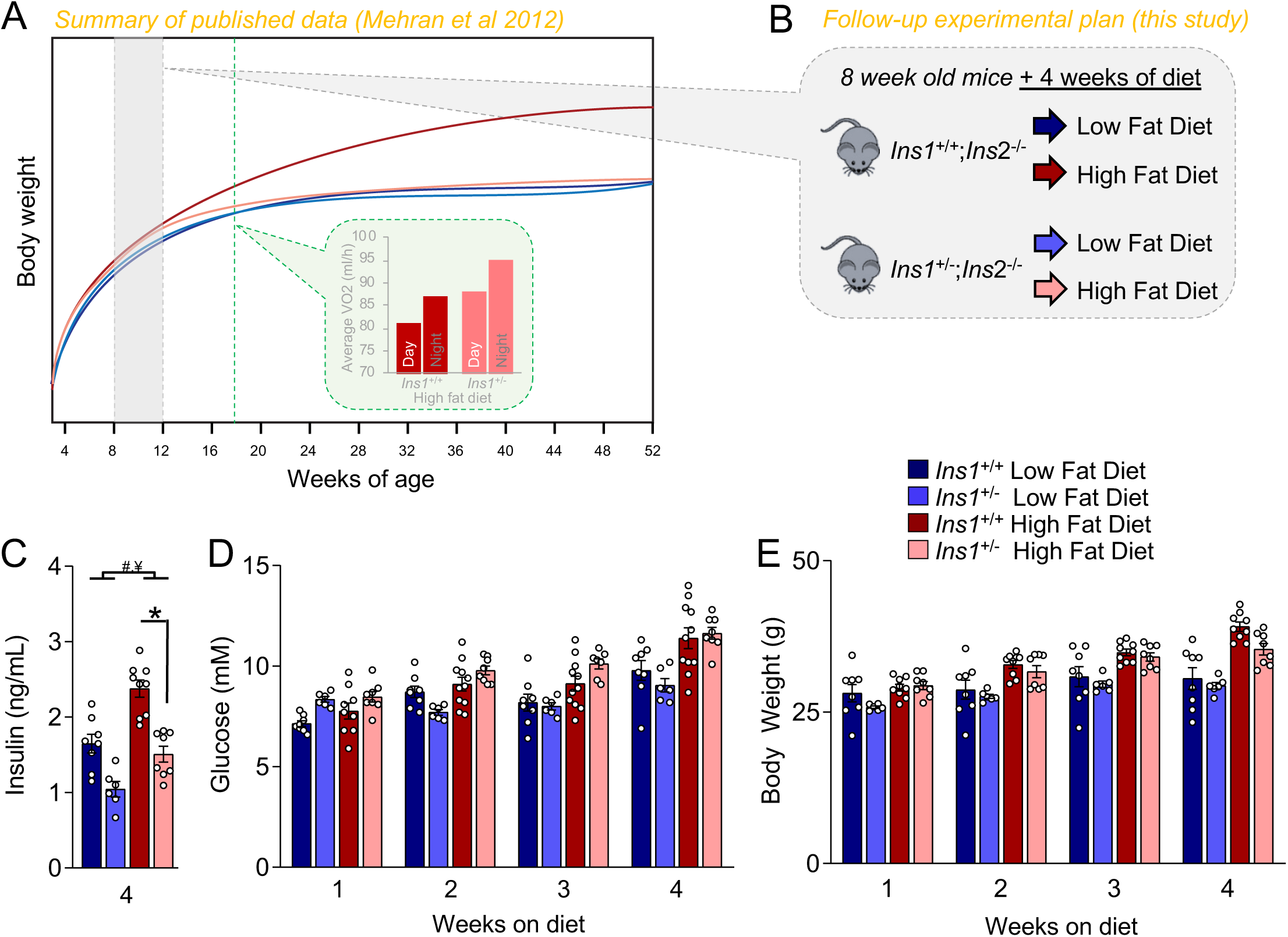
Effects of diet and *Ins1* gene dosage on circulating insulin, glucose, and body weight. **(A)** Previous data published by our group showing the consequences of long-term HFD in *Ins1*^+/-^ animals. **(B)** Experiment design showing how 8 week-old animals were randomly allocated in low fat diet or high fat diet during 4 weeks. **(C)** Insulin concentration (ng/ml) at the end of study. **(D)** Weekly glucose (mM). **(E)** Weekly body weight measurements (g). Data are presented as means ± SEM, with n=6-11. ^#^ indicates a significant diet effect, LFD vs HFD; ^¥^ indicates a significant genotype effect *Ins1*^+/+^ *vs Ins1*^+/-^. * indicates differences between individual groups.

### Effects of reduced insulin on mitochondrial oxidative complex protein in liver and muscle

Liver and muscle contribute for nearly half of energy daily expenditure in mammals (4). To investigate the effects of manipulating circulating insulin on the abundance of oxidative mitochondrial complex proteins, we conducted Western blot analysis on the central liver lobe and the gastrocnemius muscle, selected to match the tissues studied in our previous research (22). After 4 weeks of dietary intervention, no diet- or genotype-driven differences could be observed in the abundances of proteins representing complex I (NADH:ubiquinone oxidoreductase, Ndufb8), complex II (succinate dehydrogenase [ubiquinone] iron-sulfur subunit Sdhb), complex III (cytochrome b-c1 complex subunit 2, Uqcrc2), or complex V (ATP synthase, Atp5a1), with the exception of a mild increase in liver Sdhb by reduced insulin in the context of a LFD (**Figure 2A,B**). Together, these data suggest that reduced circulating insulin does not robustly affect mitochondrial oxidative metabolism proteins in liver or muscle at 12 weeks of age.

**Figure 2.**
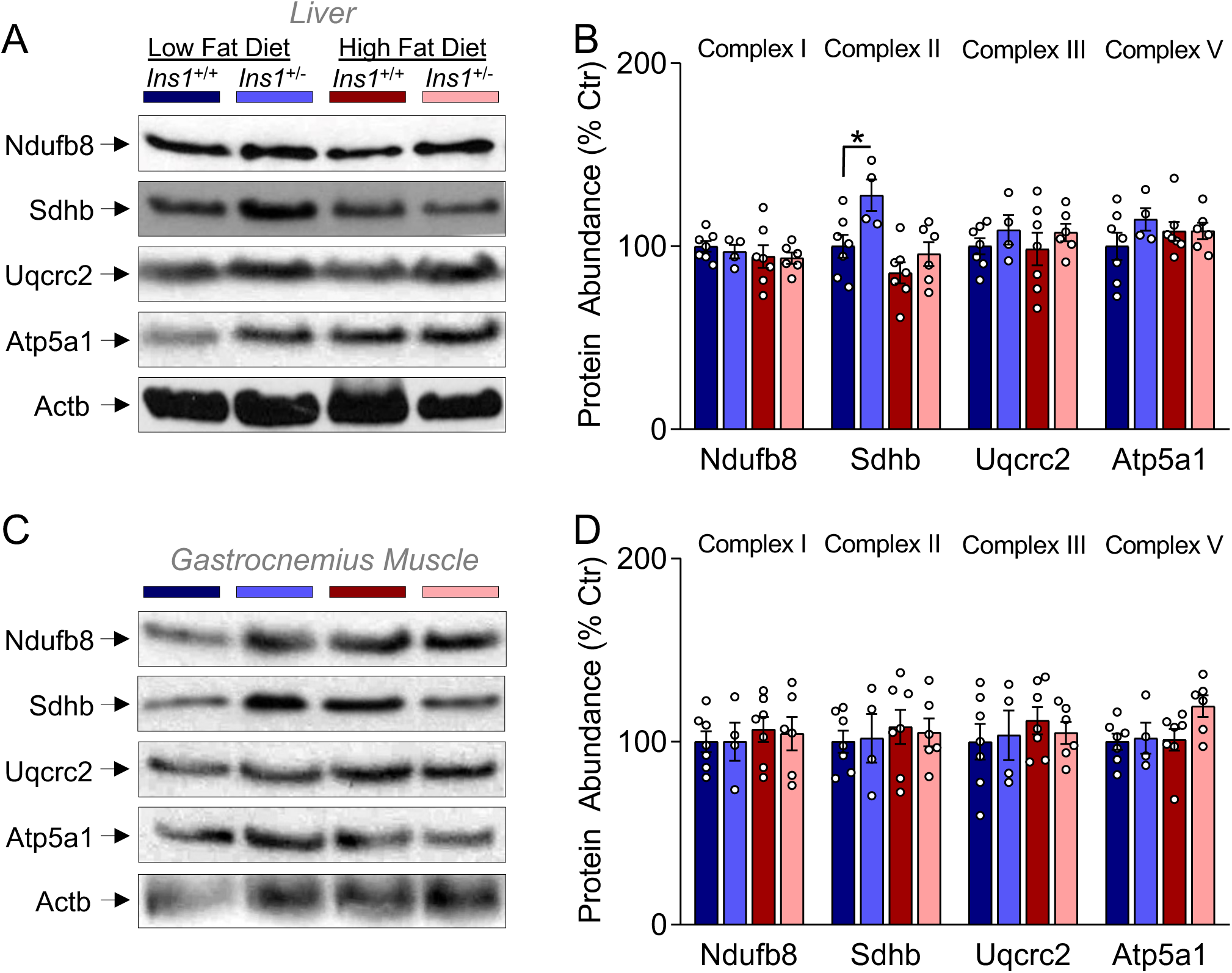
Effects of reduced insulin on mitochondrial oxidative complex protein in liver and muscle. **(A,B)** Representative western blots and associated quantification of the expression of oxidative phosphorylation complexes Nudfb8, Sdhb, Uqcrc2 and Atp5a1 in liver. **(C,D)** Representative western blots and the associated quantification of the expression of oxidative phosphorylation complexes Nudfb8, Sdhb, Uqcrc2 and Atp5a1 in skeletal muscle. Data are normalized to LFD *Ins1*^+/+^ and data are presented as mean ± SEM, with n=4-6. * indicates differences between groups.

### Effects of insulin on oxidative and uncoupling proteins in mesenteric WAT

We investigated the effects of genetic insulin reduction on oxidative mitochondrial complexes expression in three adipose tissues. In visceral fat, specifically mesenteric WAT, we found that Nudf88 abundance was paradoxically downregulated in *Ins1*^+/-^ mice compared to *Ins1*^+/+^ littermate controls (**Figure 3A,B**). Animals fed a HFD also had lower abundance of Sdhb and this was further reduced in *Ins1*^+/-^ mice (**Figure 3A,B**). HFD decreased Ucp1 expression in this fat depot and this effect was prevented in *Ins1*^+/-^ mice (**Figure 3C**). To evaluate the molecular mechanisms responsible for changes in oxidative complexes and Ucp1, we examined the abundance of Pparg, a master regulator of adipose differentiation. HFD strongly downregulated Pparg protein levels in the mesenteric independent of *Ins1* gene dosage (**Figure 3D**). This implies that the effects of reduced insulin production on Ucp1 protein levels in mesenteric WAT are independent of Pparg protein abundance. Collectively, these data point to the possibility that reducing insulin might reduce energy expenditure in mesenteric WAT.

**Figure 3.**
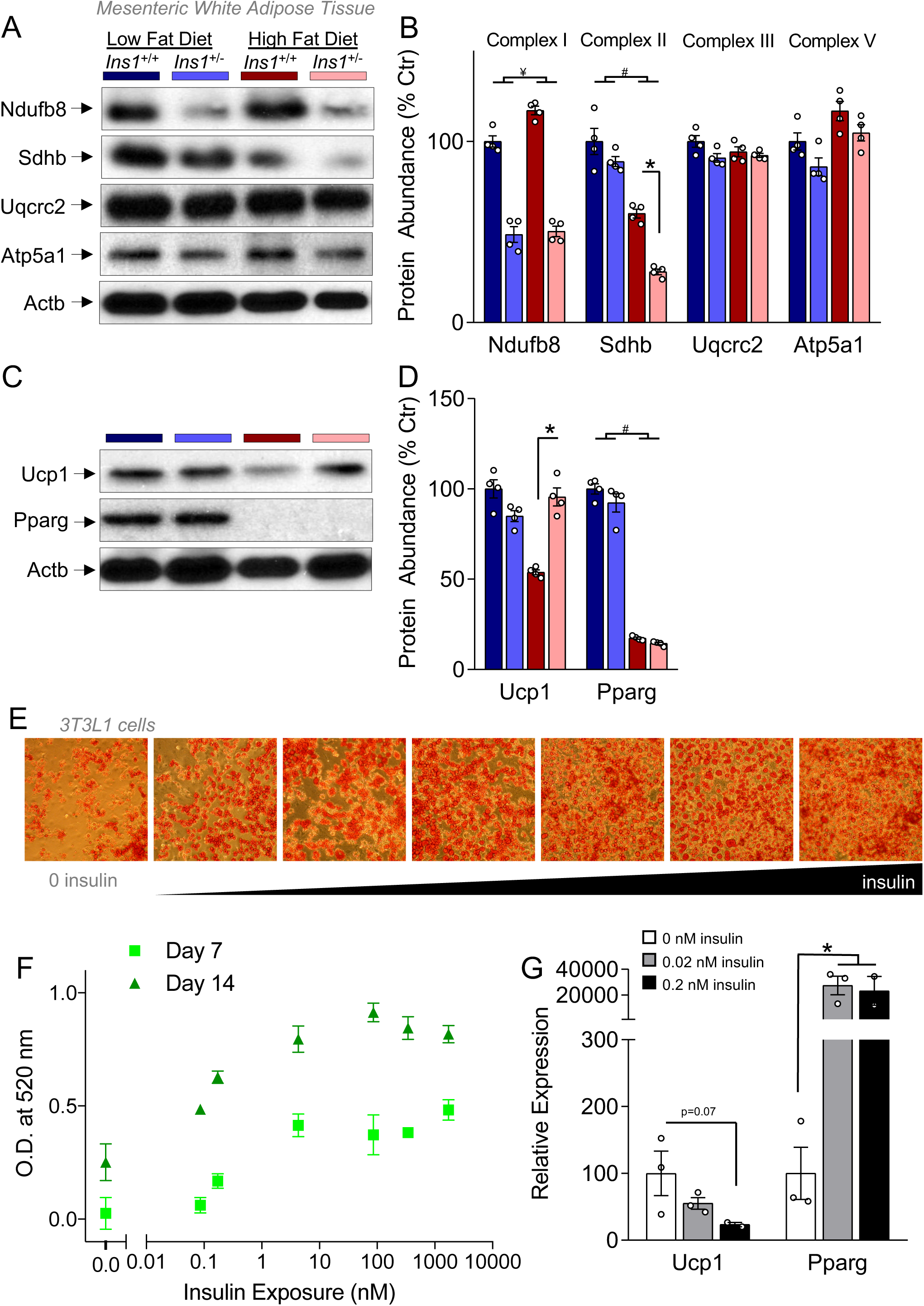
Reduced insulin selectively affects oxidative proteins expression in mesenteric white adipose tissue. **(A,B**) Representative western blots and associated quantification of the expression of oxidative Nudfb8, Sdhb, Uqcrc2 and Atp5a1, Ucp1 and Pparg in mesenteric white adipose tissue. **(C,D)** Protein abundance of Ucp1 and Pparg. Data are normalized to LFD *Ins1*^+/+^. Data are presented as means ± SEM with n=4-6. Data shown are from the same blots, where the membranes were stripped/restored between the antibody detections. ^#^ indicates a significant diet effect, LFD vs HFD; ^¥^ indicates a significant genotype effect *Ins1*^+/+^ *vs Ins1*^+/-^. * Represents differences between groups. **(E,F)** Quantification of oil red O staining in 3T3L1 cells under different insulin concentrations at 7 and 14 days of differentiation culture. Images in E are from 14 days. Data are representative of 3 independent experiments. **(G)** RNA expression at 14 days of *Ucp1* and *Pparg* in 3T3L1 cells exposed to 0, 0.02 nM and 0.2 nM of insulin in the media. Data are presented as means ± SEM. n=3. * indicates differences between individual groups.

### Insulin exposure increases 3T3L1 differentiation and Pparg expression

In order to test whether our *in vivo* results could be explained in part by direct effects on adipocytes, we evaluated the effects of multiple insulin concentrations on a cell culture preadipocyte model. After confluence, we added differentiating medium containing a range of insulin concentrations [0-10,000 nM]. Indeed, insulin had a dose-dependent effect to increase lipid accumulation at both the 7- and 14-day time points (**Figure 3E-G**). Next, we evaluated the effects of insulin doses in the physiological range on *Ucp1* and *Pparg* mRNA expression. Although we did not reach the statistical significance (p=0.07), 3T3L1 cells incubated with 0.2 nM of insulin had a reduction in *Ucp1* expression. Moreover, insulin administration was a potent activator of Pparg mRNA expression (200x) in 3T3L1 exposed to medium containing this hormone (0.02 and 0.2 nM). Together with the *in vivo* described above and previously (24), these *in vitro* studies point to the possibility that insulin can directly act on adipocytes to decrease Ucp1.

### Effects of insulin on oxidative and uncoupling proteins in inguinal WAT

Next we examined the subcutaneous inguinal WAT depot. We found that reducing insulin caused a robust increase in all mitochondrial complex proteins we assayed, across both diets (**Figure 4A,B**). There was also a stimulatory effect of HFD on Nudfb8, Sdhb and Uqcrc2 in *Ins1*^+/-^ animals (**Figure 4A,B**). Analysis of oxygen consumption rate (OCR) of isolated fat tissue using the Seahorse Bioanalyzer showed a trend towards increase OCR in *Ins1*^+/-^, but did not reach statistical significance (**Figure 4C**). These data demonstrate that reducing insulin affects visceral and subcutaneous fat pads differentially, with the later having a protein expression pattern that could increase energy expenditure. We did not find differences in Ucp1, Pparg or Prdm16:Pparg association in the inguinal WAT (**Figures 4D-G**). Sirtuins are known to regulate the cellular metabolism adipose tissue fate (28). HFD was associated with an increase in nuclear Sirt1 protein, but there was no difference in total Sirt3 abundance (**Figure 4H**). Thus, while reducing circulating insulin stimulates the expression all oxidative phosphorylation proteins in inguinal WAT, the molecule mechanisms appear to be independent of Pparg, Prdm16, Sirt1, and Sirt3 protein levels.

**Figure 4.**
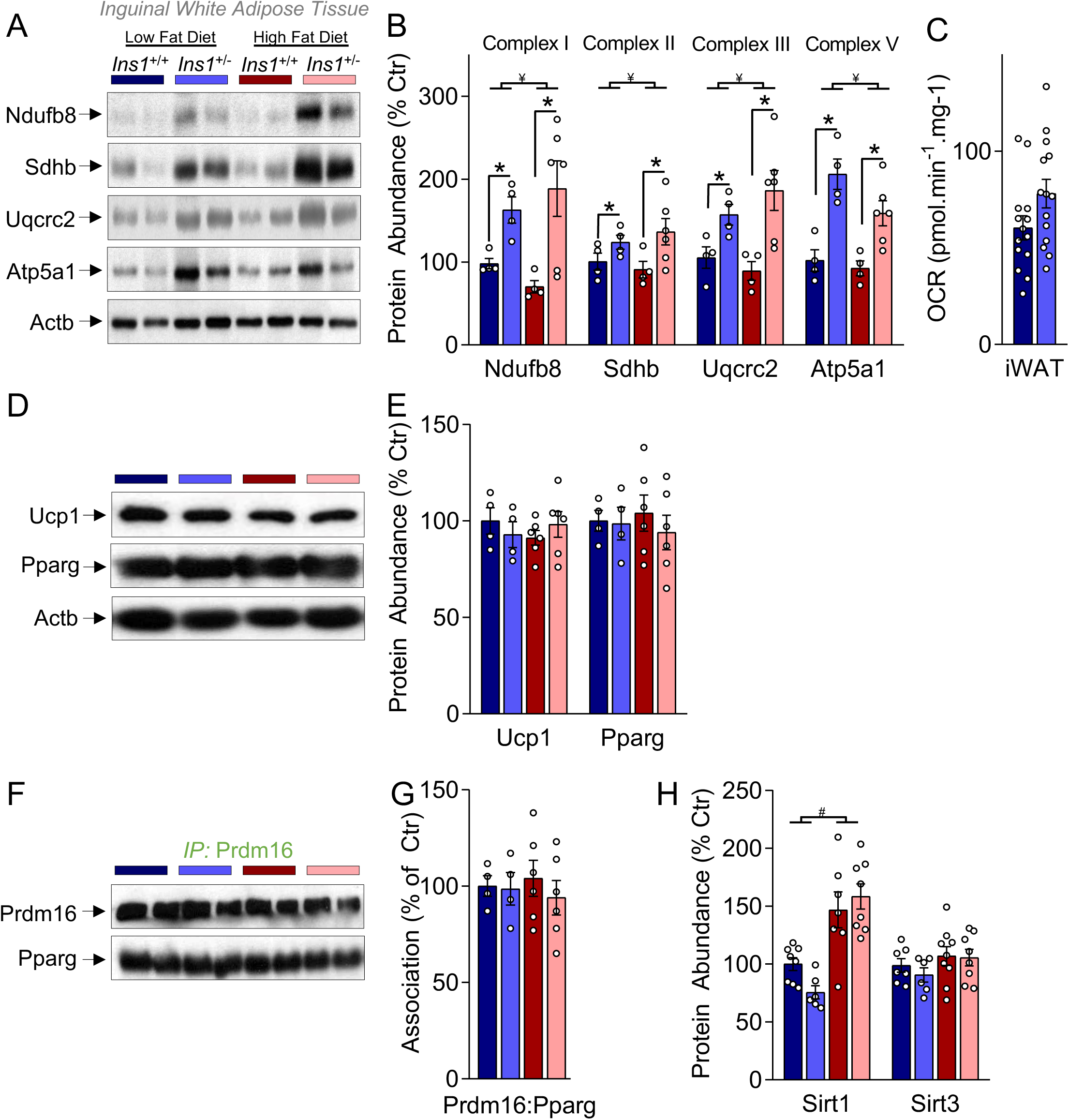
Reduced insulin increases oxidative proteins expression in inguinal white adipose tissue. **(A,B)** Representative western blots and associated quantification of the expression of oxidative Nudfb8, Sdhb, Uqcrc2 and Atp5a1 in inguinal white adipose tissue. **(C)** Basal oxygen consumption rate during basal state are normalized to LFD *Ins1*^+/+^. **(D,E)** Protein abundance of Ucp1 and Pparg. **(F,G)** Co-immunoprecipitation analysis of the Pparg:Prdm16 interaction. **(H)** Protein abundance of Sirt1 and Sirt3. Data are normalized to LFD *Ins1*^+/+^. Data are presented as means ± SEM with n=4-6. ^#^ indicates a significant diet effect, LFD vs HFD; ^¥^ indicates a significant genotype effect *Ins1*^+/+^ *vs Ins1*^+/-^. * indicates differences between individual groups.

### Effects of reduced insulin on oxidative protein expression in BAT

We also examined the effects of reduced insulin levels on the same set of proteins in interscapular BAT. HFD feeding reduced the abundance of Sdhb protein (LFD vs HFD; p=0.01), although this effect was partially prevented in *Ins1*^+/-^ mice (**Figure 5B**). In BAT, HFD reduced the abundance of ATP synthase (p=0.05), and that was exacerbated by reducing insulin production (**Figure 5B**). We found no statistical differences in OCR (p=0.07) between LFD *Ins1*^+/-^ and *Ins1*^+/+^ mice (not shown). Ucp1 protein abundance and Prdm16:Pparg association were increase by HFD, and this effect was potentiated in the context of reduced insulin (**Figure 5C-F**). Interestingly, Pparg abundance was reduced in these cells (**Figure 5C,D**). We evaluated the Sirt1 and Sirt3 activity and protein content in BAT, and found that Sirt1 activity was higher in the BAT of HFD *Ins1*^+/-^ compared to HFD *Ins1*^+/+^ in all time points, despite a lack of differences in nuclear Sirt1 content (**Figure 5G,H**). We observed a HFD-dependent increase in the Sirt3 (mitochondrial) activity and Sirt3 content in BAT (**Figures 5G,H)**. These data demonstrate that, in the context of HFD, reduced insulin increases Ucp1 protein levels, potentially through a mechanism involving the association of Prdm16 with Pparg and the upregulation of Sirt3.

**Figure 5.**
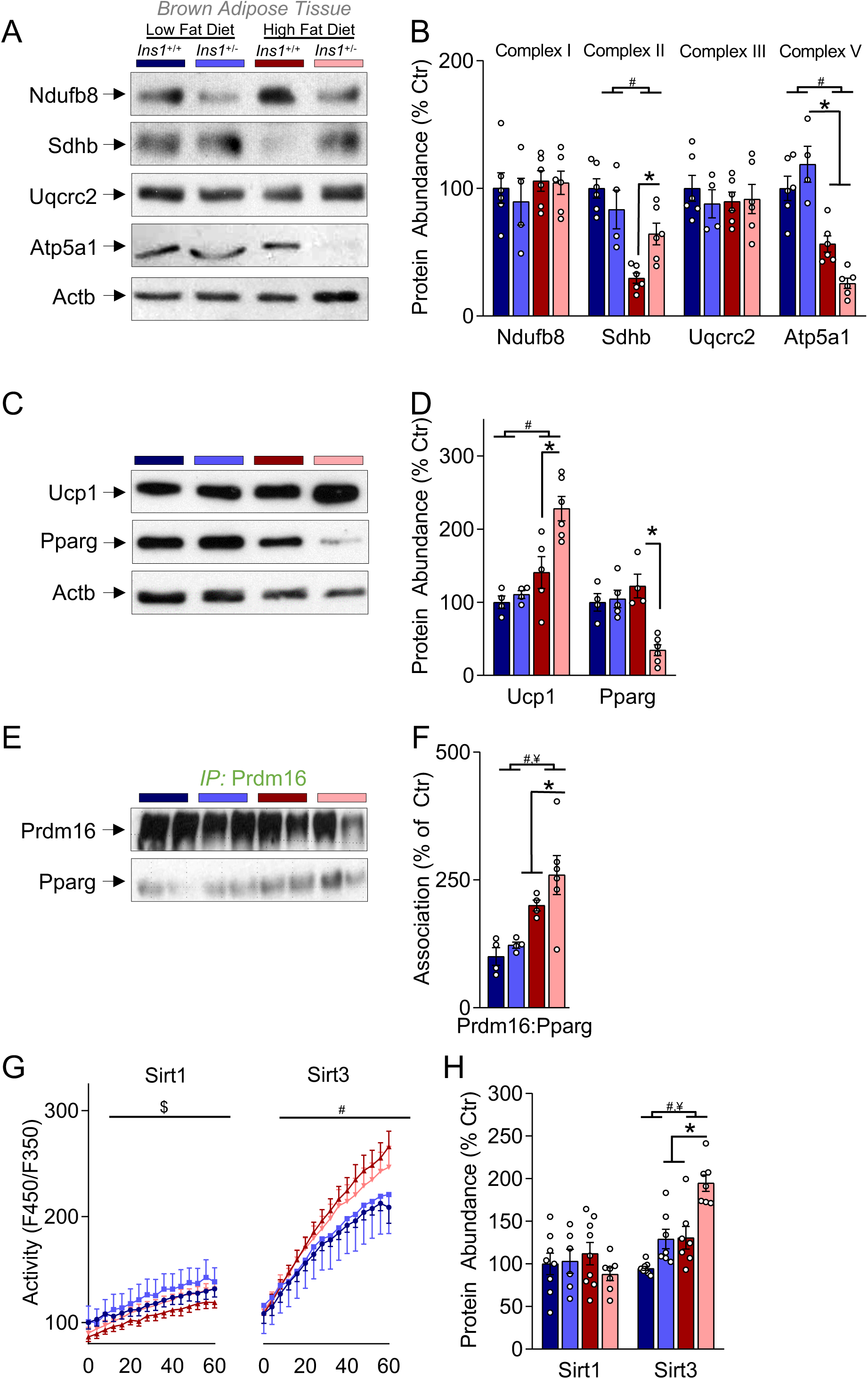
Reduced insulin increases Ucp1 protein and Prdm16:Pparg interaction in BAT. **(A,B)** Representative western blots and associated quantification of the abundance of oxidative Nudfb8, Sdhb, Uqcrc2 and Atp5a1 proteins. **(C,D)** Protein abundance of Ucp1 and Pparg. **(E,F)** Immunoprecipitation analysis was used to detect Pparg:Prdm16 interaction. **(G,H)** Protein abundance and activity of Sirt1 and Sirt3. Data are normalized to LFD *Ins1*^+/+^. Data are presented as means ± SEM with n=4-6. Data shown are from the same blots (2 biological replicates per blot), where the membranes were stripped/restored between the antibody detections. Immunoblots in panels A and C were probed in the order presented, including a reprobing of Actb. Data are presented as means ± SEM with n=4-6. ^#^ indicates a significant diet effect, LFD vs HFD; ^¥^ indicates a significant genotype effect *Ins1*^+/+^ *vs Ins1*^+/-. $^ Difference between HFD *Ins1*^+/+^ vs HFD *Ins1*^+/-^. * indicates differences between individual groups.

## Discussion

In the present study, we tested the hypothesis that lowering insulin levels might trigger metabolic reprogramming in various tissues to increase energy expenditure and protect mice against HFD-induced obesity (22). We studied mice that are genetically incapable of hyperinsulinemia at a young age before effects on body weight and blood glucose are apparent (22). We found that after 4 weeks of HFD diet, 12 week-old mice with genetically reduced insulin had reprogrammed their mesenteric WAT, inguinal WAT, and BAT in distinct ways, while skeletal muscle and liver were largely unaffected at this time point. Specifically, mesenteric WAT and interscapular BAT upregulated Ucp1 protein levels, whereas inguinal WAT increased the abundance of key oxidative phosphorylation proteins. Although mechanistically different, both forms of adipose tissue metabolic remodeling would be expected to favour the increased energy expenditure we previously observed experimentally at 18 weeks in this animal model (22).

Insulin is the body’s principle diet-regulated anabolic hormone and it acts directly on adipose tissue. Adipose tissue selective insulin receptor knockout (FIRKO) animals are protected against diet-induced obesity and glucose intolerance (3). In our previous studies, we have shown that genetically suppressing hyperinsulinemia is sufficient to prevent diet-induced obesity. In our first of a series of studies, we found that preventing hyperinsulinemia increased energy expenditure and we examined mRNA expression of known metabolism regulating genes in muscle, liver, epididymal WAT, and BAT of 1-year old mice to glean information about the underlying molecular mechanisms. These studies uncovered elevated *Ucp1* mRNA expression and protein levels in epididymal WAT in mice with reduced insulin, suggesting that hyperinsulinemia normally suppresses Ucp1 abundance in WAT (22). Although *Ucp1* gene expression in BAT was unaffected (22), we did observe an increase in BAT weight in subsequent cohorts (N.M. Templeman, unpublished observations), which likely contributed to the increased energy expenditure in this model. Subsequent studies used global transcriptomics and proteomics in liver and muscle to examine mice with a different configuration of germline insulin gene mutations that were also protected from diet-induced obesity (39) and age-dependent insulin resistance (38). Consistent with the present results, we did not observe a robust signature supporting increased energy expenditure in muscle either mouse model of constitutively reduced insulin (22, 37, 38). These results also corroborate other work showing that short-term HFD feeding does not robustly alter insulin sensitivity in liver (15) or skeletal muscle (33). Using an inducible model of partial insulin gene deletion in obese 16 week old mice, we found suppressed insulin production for 5 weeks led to reduced mass of gonadal and perirenal fat pads, but not inguinal WAT, mesenteric WAT or BAT (25)(27). *Ucp1* was not different in the gonadal fat pads of these 19 week-old mice when examined using RNAseq. Thus, until this present study the effects of reduced hyperinsulinemia on energy expenditure mechanisms not been examined young mice.

Here, we report multiple, tissue-specific mechanisms for upregulating energy expenditure in mice with genetically suppressed hyperinsulinemia. One mechanism, which we have previously identified in these *Ins1*^+/-^;*Ins2*^-/-^ mice, is the upregulation of mitochondrial Ucp1 protein levels. Upregulation of Ucp1, a principle component of so-called WAT ‘browning’ is a well-established mechanism to increase energy expenditure in WAT by ‘wasting’ mitochondrial proton potential as heat. In the young mice we studied, the upregulation of Ucp1 was specific to mesenteric WAT and BAT. We provided evidence from 3T3L cultured adipocytes that insulin reduces Ucp1 expression cell autonomously, consistent with *in vitro* work from other groups (16) and the upregulation of Ucp1 by reduced insulin we observed *in vivo*. The upregulation of Ucp1 may have additional consequences. For example, our data on mesenteric WAT and BAT lead us to speculate that increased Ucp1 abundance may reduce the demand on ATP synthase (Atp5a1), without changing mitochondrial content. Indeed, BAT from *Ucp1*^-/-^ mice has increased ATP synthase membrane subunit C locus 1 (*Atp5g1*) and increased ROS production, as well as mitochondrial electron transport chain dysfunction (14). Collectively, these data point to the upregulation of Ucp1 protein levels as a mechanism for increasing energy expenditure in some, but not all, fat depots.

To further assess the molecular mechanisms involved, we measured the protein abundance of the master adipogenesis regulator Pparg in 3 adipose depots. In mesenteric WAT and BAT where Ucp1 was increased, we found a 60% reduction in *Pparg* expression under the same conditions of HFD with reduced hyperinsulinemia. We have previously found in that 1 year old HFD-fed *Ins1*^+/-^ mice had a ∼40% reduction in *Pparg* mRNA in inguinal WAT at 1 year of age (22). *Pparg* deletion in adipose tissue is sufficient to protect mice against HFD-induced obesity (12). Activating Ppar promotes WAT browning via Prdm16 (24). In BAT, Ucp1 upregulation was associated with an increase in the association of Prdm16 with Pparg, while this immunoprecipitation was not successful in mesenteric WAT in our hands.

Sirtuins are multifunctional enzymes that have multiple roles in adipocyte metabolism (11, 36). In particular, the nuclear localized Sirt1 protein upregulates Ucp1 and browning of WAT through Pparg (28). Sirt1 can also deacetylate Sirt3 (8, 9, 17), which in turn can directly increase *Ucp1* expression (32). *Sirt3* gene expression is controlled by Pparg1ca (8, 17). We were not able to measure the activity of Sirt1 and Sirt3 enzymes in WAT, due to low expression. In BAT, active Sirt1 deacetylates two Pparg residues, which promotes interaction with Prdm16 (28). This complex is responsible to increase the transcription of the Sirt3. Sirt3 regulates mitochondrial biogenesis, oxygen consumption, and ROS production (8, 17), shifting cells to more efficient energy production with lower ROS and higher aerobic output (26). In our studies, BAT from HFD-fed *Ins1*^+/-^;*Ins2*^-/-^ mice had increased binding of Prdm16 to Pparg, increased Sirt1 activity, increased Sirt3 abundance, and increased Ucp1 abundance. We were unable to observe statistically significant differences in Sirt3 activity between genotypes, but a prominent effect of HFD was found. The adaptation we observed are consistent with the idea that lowering insulin levels can preserve the cellular metabolic fitness. Since BAT plays a larger role than WAT in total energy expenditure in mice housed at room temperature (14), these changes in Ucp1 could be physiologically significant, along with an increase in BAT mass.

Upregulation of oxidative phosphorylation proteins is also a mechanism by which adipose tissue can increase energy expenditure in fat-specific insulin receptor knockout animals (13). Specifically, we found that in inguinal WAT, preventing hyperinsulinemia was sufficient to induce a 1.5-to 2-fold increase in the protein abundance of key mitochondrial oxidative phosphorylation component across both diets tested. Other studies have found that insulin signaling downregulates mitochondrial oxidative phosphorylation genes in liver and skeletal muscle via Akt inhibition of Ppargc1a (20, 35). At the young age we studied, we did not observe effects in liver or muscle, other than reduced liver Sdhb expression the context of HFD. In contrast to the effects in inguinal WAT, mesenteric WAT exhibited reduced Ndufb8 and Sdhb in *Ins1*^*+/-*^ animals. In BAT, HFD reduced Atp5a1 and Sdhb abundance, although the later effect was counteracted by preventing hyperinsulinemia. Reductions in Sdhb are associated to insulin resistance and visceral adipogenesis in humans (23). Further studies are necessary to elucidate the mechanism behind the differential regulation of oxidative phosphorylation complex components in inguinal WAT, mesenteric WAT, and BAT.

Mitochondrial dysfunction is considered to be one of the major outcomes of hyperinsulinemia, insulin resistance and obesity (5, 21, 29). HFD can overwhelm mitochondrial biogenesis in WAT and decreases WAT energy expenditure (30). WAT mitochondrial loss has been demonstrated as a strong predictor of type 2 diabetes (41). Nudfb8 expression has the strongest association with mitochondrial content (18). Whether reduction of circulating insulin in our model increased mitochondrial mass and upregulated specific proteins in a similar number of mitochondria, or a combination of effects, is not clear and would require detained histology.

Our study had several limitations that will have to be resolved with future work. For example, there are other possible mechanisms by which mice can up-regulate energy expenditure in the context of reduced circulating insulin, which we did not directly address in this study. Although we did not find significant physical activity changes in our previous of *Ins1*^+/-^;*Ins2*^-/-^ studies, there was a trend towards increased physical activity that could be explored in larger cohorts. Housing temperature was set below thermoneutrality, and this is known to affects the relative importance of changes in Ucp1 (9). Although we did not evaluate female animals, previous studies from our group showed similar body weight results in Ins1^+/-^ animals (7, 43). The current study aimed to investigate the mechanisms underlying a specific set of previous findings and it was for this reason that we decided to evaluate the same tissues as before, and exclude other muscle types, for example. Ideally, we would have also obtained tissue-level energy expenditure measurements in this study, but our efforts to examine oxygen consumption in several tissues were unsuccessful. We were also unable to examine some key mitochondrial proteins and protein-protein interactions in WAT, due to the lack of their abundance. However, despite these limitations, we feel our study contributes to the understanding of how the prevention of hyperinsulinemia can increase energy expenditure and protect mice from weight gain.

In conclusion, it is known that WAT has a key role in development of insulin resistance and obesity, and it’s endocrine function affects virtually all cells (10). Our study provides evidence that adipose tissue depots remodel their energy expenditure capabilities using distinct mechanisms. Collectively, we provide the rationale for modestly reducing insulin levels, perhaps via lifestyle changes, to exert beneficial effects on adipose tissue metabolism by reprogramming its efficiency towards increased energy expenditure.

## Author Contributions

JDB co-conceived and designed the study, performed most of the experiments and wrote the manuscript. PO conducted tissue extraction and collaborated on the Seahorse experiments. LL was responsible for part of the western blotting experiments. OH, NMT, and GEL designed and conducted the 3T3L1 experiments. SW performed statistical analysis and edited the manuscript. GEL edited the manuscript. JDJ co-conceived and designed the study, edited the manuscript and is the guarantor of this work.

## Funding

This research was funded by a CIHR Project grant PJT168857 to JDJ. JDB was supported by a travel/salary grant from the São Paulo Research Foundation (FAPESP-2016/23251-7). GEL holds the Canada Research Chair in Adipocyte Development.

## Notes

### Competing Interest Statement

The authors have declared no competing interest.

